# PLANT-Dx: A Molecular Diagnostic for Point of Use Detection of Plant Pathogens

**DOI:** 10.1101/498998

**Authors:** M. Verosloff, J. Chappell, K. L. Perry, J. R. Thompson, J. B. Lucks

## Abstract

Synthetic biology based diagnostic technologies have improved upon gold standard diagnostic methodologies by decreasing cost, increasing accuracy, and enhancing portability. However there has been little effort in adapting these technologies towards applications related to point-of-use monitoring of plant and crop health. Here, we take a step towards this vision by developing an approach that couples isothermal amplification of specific plant pathogen genomic sequences with customizable synthetic RNA regulators that are designed to trigger the production of a colorimetric output in cell-free gene expression reactions. We demonstrate our system can sense viral derived sequences with high-sensitivity and specificity, and can be utilized to directly detect viruses from infected plant material. Furthermore, we demonstrate that the entire system can operate using only body heat and naked-eye visual analysis of outputs. We anticipate these strategies to be important components of user-friendly and deployable diagnostic systems that can be configured to detect a range of important plant pathogens.

---

Synthetic biology has recently contributed to multiple advances in point-of-use nucleic acid diagnostic (PoUD) technologies (1). These technologies leverage isothermal strategies to amplify target nucleic acids, with new advances in detection of these targets using strand-displacement (2), or CRISPR-based methods (3, 4) that produce fluorescent readouts, or RNA toehold switches that control translation of enzymes that produce colored compounds (5). Overall these technologies can be used for sensitive detection of pathogen-derived nucleic acids in complex matrices in field-deployable formats that significantly improve upon current laboratory-intensive PCR-based approaches.

To date, most synthetic biology diagnostic efforts have focused on detecting pathogens that impact human health. However, there is great potential to leverage these technologies for detecting plant pathogens. In the United States alone, plant pathogens account for an estimated $33 billion annual loss in agricultural productivity (6). Worldwide, losses in crop yields due to plant pathogens can be more severe and contribute to food scarcity and famine (7). Among plant pathogens, wide host range viral species such as cucumber mosaic virus (CMV) and potato virus Y (PVY) are particularly devastating as they infect hundreds of plant species, including agriculturally important species such as beans, maize, and potatoes (8). PoUDs are an important component of strategies to combat the impacts of these pathogens, as timely identification can lead to the rapid deployment of methods for mitigation and containment. However, current plant pathogen PoUD strategies use a range of approaches including antibody-based detection which lacks sensitivity (9), or isothermal amplification that by itself does not generate convenient visual outputs that are amenable to field use.

To address these shortcomings, we sought to develop a PoUD system called PLANT-Dx (**P**oint of use **LA**boratory for **N**ucleic acids in a **T**ube) that combines the sensitivity of isothermal strategies to amplify target plant pathogen nucleic acids (10), with the designability of synthetic gene regulatory systems (11) and robustness of cell free gene expression reactions (12) to produce colorimetric outputs that are visible to the naked-eye (Figure 1A). The overall approach of our PLANT-Dx system is to convert plant pathogen nucleic acids into constructs encoding designed synthetic RNA regulators that when produced activate an RNA genetic switch controlling the expression of an enzyme that catalyzes a color change. The RNA genetic switch thus serves a role as a signal processing layer that filters the noisy output of isothermal amplification products (10), only triggering the production of color for correctly amplified on-target viral sequences.

**Figure 1:**
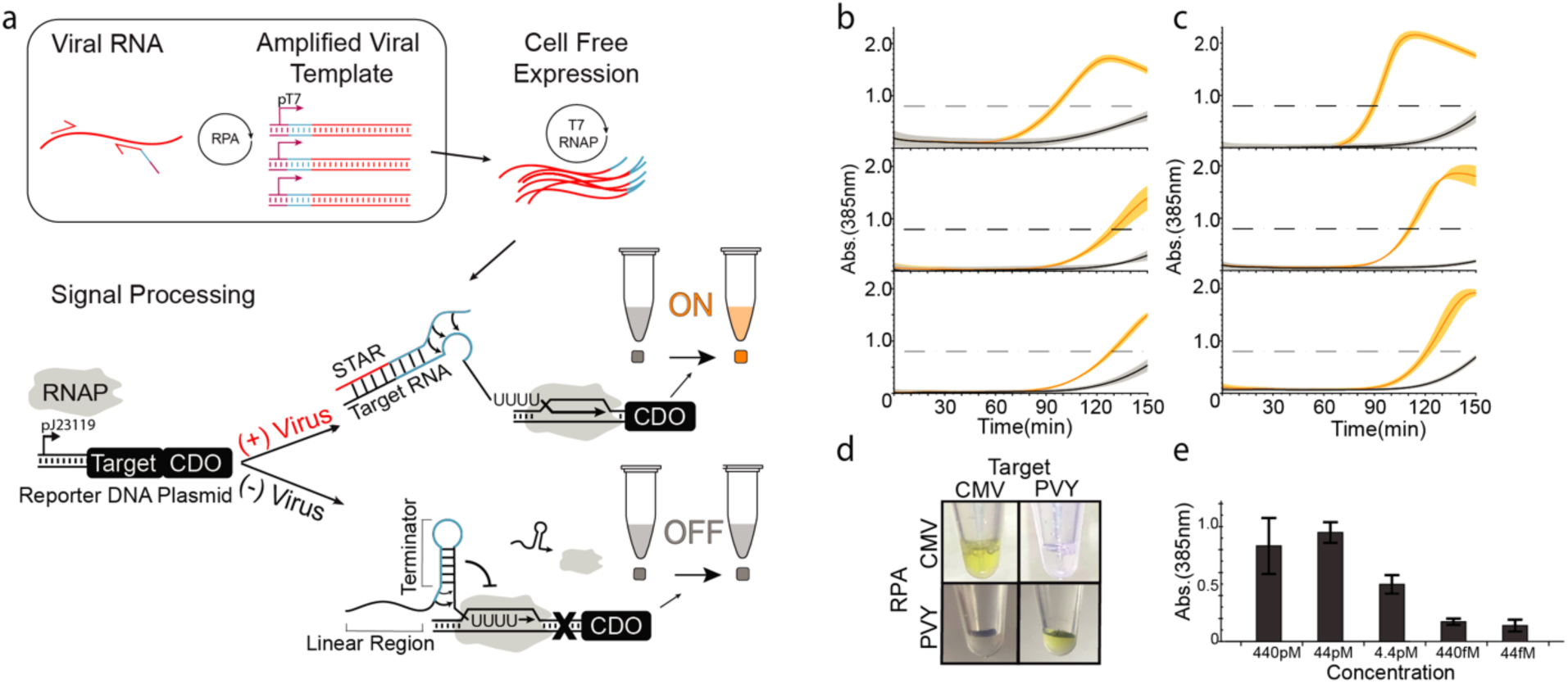
Basic operation of PLANT-Dx. (a) Schematic overview. Viral RNA is amplified by recombinase polymerase amplification (RPA) into DNA templates that contain a T7 polymerase promoter (purple), a portion of a small transcription activating RNA (STAR) sequence (blue), and a portion of the viral RNA sequence (red). Cell free expression of these templates produces viral sequence-derived STAR RNA which triggers the production of catechol 2,3-dioxygenase (CDO), which in turn converts hydroxymuconic semialdehyde into a yellow colored compound which is visible by eye near a 385nm absorbance value of 0.8 (dashed line, SI Figure 3, 4). (b) Demonstrated ability to detect cucumber mosaic virus (CMV) sequences from in vitro transcription (IVT) RNA products (orange) versus control (grey) samples, by 120 minutes across three different days. (c) Demonstrated ability to detect potato virus Y (PVY) based IVT RNA (orange) versus control (grey) samples by 120 minutes across three different days. (d) An orthogonality matrix of cell-free reactions challenging RPA products from different IVT sources against different STAR-Target-CDO constructs showing positive results (yellow) only for cognate combinations at 150 minutes. (e) Serial dilution of CMV IVT RNA was used to determine a limit of detection in between 4.4pM and 440fM after 150 minutes of reaction. Data in (b), (c) represents mean values and error bars represent s.d. of n = 3 technical replicates, and (e) represents mean values and error bars represent s.d. of n = 3 biological replicates that each had n = 3 technical replicates (n=9 total).

PLANT-Dx works by first using recombinase polymerase amplification (RPA) (13) (Supplementary Figure 1) to amplify a target region of a plant pathogen genome to produce a double-stranded DNA construct that encodes the synthesis of a synthetic RNA regulator called a Small Transcription Activating RNA (STAR) (SI Figure 1) (11). These DNA templates are then used to direct the transcription of STARs within a cell-free gene expression reaction (12), which when produced, activates the transcription of a STAR-regulated construct encoding the enzyme catechol 2,3-dioxygenase (CDO) (14) (Figure 1A). Only when the pathogen is present is the RPA product made, leading to expression of CDO, which in turn converts the colorless catechol compound into a visible yellow product. Here we show that this design can detect CMV in infected plant lysate with a low picomolar sensitivity, and can be configured to detect nucleic acids from different viral genomes without crosstalk. In addition, we show that this design requires only simple mixing and body heat to induce a color change, which we anticipate will facilitate deployment to field settings.

## Results

To develop PLANT-Dx, we first sought to create pathogen detecting molecular sensors based upon the Small Transcription Activating RNA (STAR) regulatory system (11). This transcription activation system is based upon conditional formation of a terminator hairpin located within a target RNA upstream of a gene to be regulated: alone, the terminator hairpin forms and halts transcription of the downstream gene, while in the presence of a specific *trans-acting* STAR the hairpin cannot form and transcription proceeds (SI Figure 2). Previous work showed that the STAR linear binding region can be changed to produce highly functional and orthogonal variants (15). Here we sought to utilize this by replacing the linear binding region with sequences derived from genomic pathogen DNA to create new viral sensors. To do this, we utilized the secondary structure prediction algorithm NUPACK to identify regions within the genomes of cucumber mosaic virus (CMV) and potato virus (PVY), that are predicted computationally to be unstructured for target RNA design (Supplementary Note 1) (16). Once viral STARs were designed, reporter DNA constructs were then created in which these target RNA sequences were placed downstream of a constitutive T7 promoter and upstream of the CDO reporter gene. We next designed RPA primer sets to amplify and transform a pathogen’s genomic material into a DNA construct capable of synthesizing a functional STAR. Specifically, a T7 promoter and anti-terminator STAR sequence were added to the 5’ end of a reverse RPA primer, which when combined with a forward primer, amplified a 40 nt viral sequence to produce a double-stranded DNA encoding the designed STAR which contained the target viral sequence. In this way, we anticipated that combining the CDO-encoding reporter construct and RPA amplified DNA into a cell-free gene expression reaction (12, 17) would lead to the production of a detectable colorimetric output signal.

**Figure 2:**
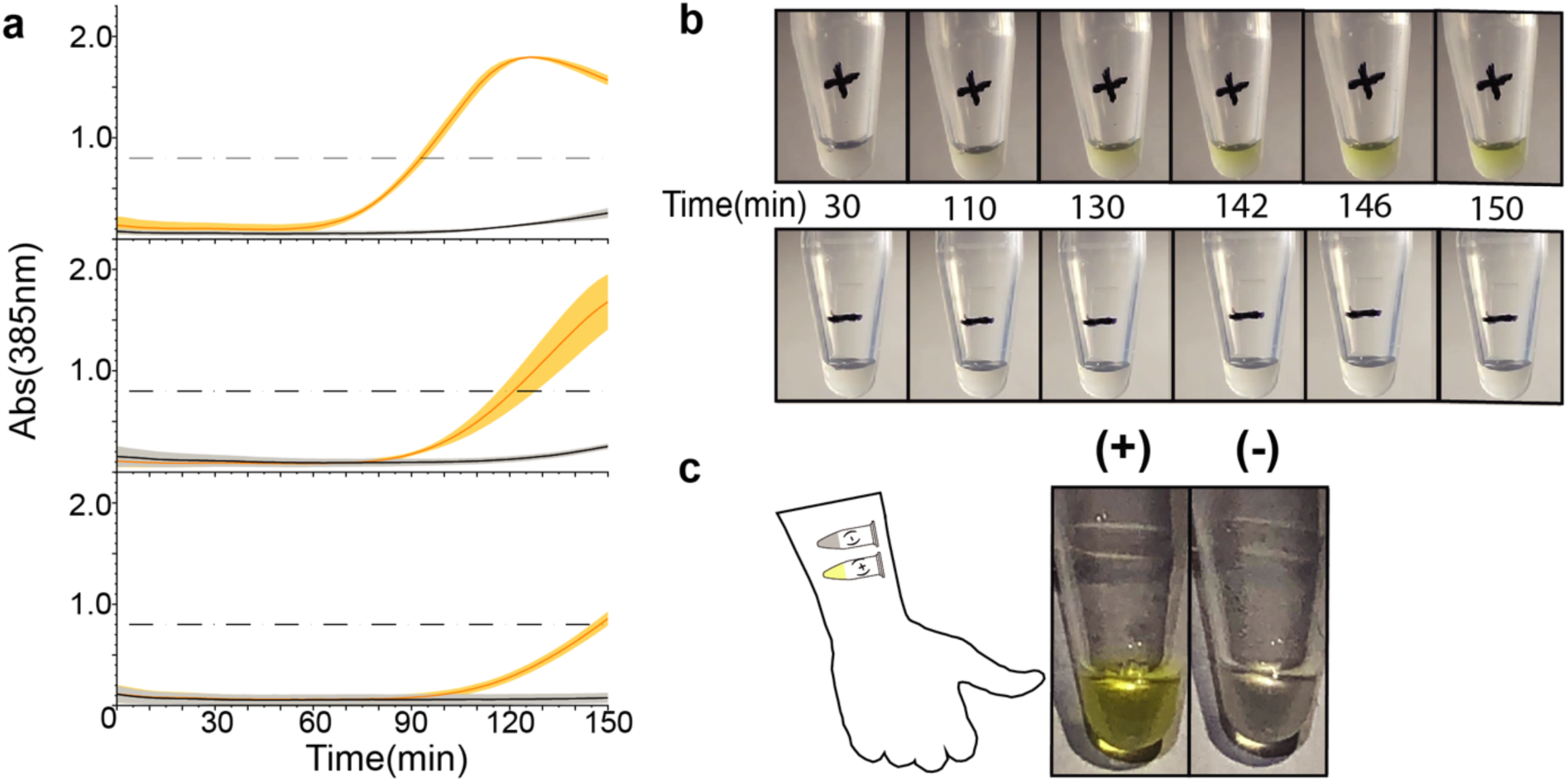
Implementing a PLANT-Dx diagnostic. (a) PLANT-Dx reactions were used to detect CMV from infected *N. tabacum* derived lysate (orange) versus uninfected plant-derived lysate (grey), across three different days. Data displayed as in Figure 1B,C. (b) Time lapse photos of cell-free reactions containing RPA products from CMV infected lysate (+) or uninfected lysate (−). (c) Demonstration that PLANT-Dx can operate using only body heat as a heat source. RPA and cell-free reactions were run by taping tubes on the inner arm for 40, and 150 minutes, respectively. A yellow color was observed for the CMV-infected lysate sample (+) and not the uninfected lysate control (−). Data in (a) represents mean values and error bars represent s.d. of n = 3 technical replicates.

We began by investigating the ability of PLANT-Dx to detect the presence of *in vitro* transcribed (IVT) RNA designed to mimic specific target regions of cucumber mosaic virus (CMV). We observed rapid color accumulation in samples containing 1 nM of purified transcription product versus the no-RNA negative control (Figure 1B). To test for modularity, we further developed sensors and primer sets for the detection of potato virus Y (PVY), and confirmed function with the same assay (Figure 1C). The specificity of our system was also tested by interrogating the crosstalk between the product of various RPA reactions and noncognate molecular sensors. Specifically, we tested color production from cell-free reactions containing the reporter DNA construct for CMV with the PVY IVT-derived RPA product, as well as the converse, and found color production only between cognate pairs of input RPA and reporter constructs (Figure 1D). We next interrogated the inherent limit of detection of our system through titration of input IVT products (Figure 1E). This demonstrated our ability to detect the presence of target nucleic acid sequences down to the picomolar range. Surprisingly, this sensitivity is lower than that previously reported for RPA (3) and is most likely due to loss in amplification efficiency from the addition of the long overhangs present within our primer sets.

We next set out to determine whether this methodology was able to differentiate between plant lysate obtained from healthy plants versus lysate infected with CMV virus. To test this, we input 1 uL of CMV-infected plant lysate, or an equivalent volume of a non-infected plant lysate control, into the PLANT-Dx reaction system. Here, we observed rapid color change only from reactions with infected lysate when compared to healthy lysate (Figure 2A). To demonstrate that this assay can be monitored by eye, reactions were carried out and filmed within a 31 °C incubator (Figure 2B). With the naked eye, we detected accumulation of a yellow color only within reactions that were incubated with infected lysate, while no such production was witnessed in reactions with uninfected lysate.

A notable drawback of current gold standard diagnostics is the need for peripheral equipment for either amplification or visualization of outputs. Even simple heating elements for controlled incubations are a major hindrance during deployment within the field and can be cost-prohibitive. We sought to exploit the flexible temperature requirements of both RPA and cell free gene expression reactions by attempting to run our diagnostic reactions for CMV infected lysate using only body heat. This resulted in clear yellow color only in the presence of infected lysate, with no major difference observed between these reactions and those previously incubated within a thermocycler and plate reader (Figure 2C).

## Discussion

Here, we have demonstrated a novel scheme for combining isothermal amplification and custom synthetic biology viral sensors for the detection of the important plant pathogen cucumber mosaic virus (CMV) from infected plant lysate. Building off of previously elucidated STAR design rules (15), we have shown that our molecular sensors can be efficiently designed, built, and implemented for use in this important plant diagnostic context. The use of STARs in PLANT-Dx complements previous uses of toehold translational switches for similar purposes in human viral diagnostics (5), and could lead to more powerful combinations of the two technologies in the future. In addition, we have shown that these reactions can be readily run without the need for extraneous heating or visualization equipment. The ability of our methodology to selectively detect genomic sequences from cucumber mosaic virus (CMV) and potato virus (PVY) highlight the ability of the growing methodologies and design principles within RNA synthetic biology (18) to contribute to real world applications. We hope that these developments can be incorporated within other synthetic biology-based diagnostic platforms (5) to enable PoUDs to be developed and delivered to regions of the world that need them most.

## Supporting information

## Author Information

M.V., J.C., K.P., J.T., and J.B.L designed the study. M.V and J.T. performed the experiments. M.V. and J.B.L performed the data analysis. All authors contributed to the preparation of this manuscript.

## Acknowledgments

We wish to acknowledge the contributions of Dr. Khalid Alam for help in ensuring reproducibility and proofreading of this manuscript, and Dr. Melissa Takahashi for inspiring the combination of RPA and STARs for pathogen detection. This work was supported by an NSF CAREER Award (1452441 to J. B. L.), Searle Funds at the Chicago Community Trust (to J. B. L.), and a Grand Challenges Explorations Grant from the Bill & Melinda Gates Foundation (OPP1199439 to J. B. L.).

