## Supplementary material for "PLANT-Dx: A Molecular Diagnostic for Point of Use Detection of Plant Pathogens"

### Material and Methods

**Template Assembly:** All DNA oligonucleotides utilized in this work were ordered from Integrated DNA Technologies. The plasmid constructs for target reporter expression were designed utilizing a NUPACK script (Supplementary Note 1). Inverse PCR was then employed to clone desired target reporter cassette constructs into a low copy p15A backbone. Assembly reactions were transformed into NEB® Turbo Competent *E. coli*, colonies isolated, and sequence confirmed (Quintara Biosciences). A table of all DNA sequences used in this work can be found below in Supplementary Table 1. Chi6 DNA stocks for cell free gene expression reactions were assembled by annealing complimentary oligonucleotides on a thermocycler programed to begin at 95$^{o}$C and drop stepwise to 25$^{o}$C at a rate of -0.1$^{o}$C/s throughout 3 minutes.

**PLANT-Dx Reactions**: PLANT-Dx consists of two reaction steps. First Small Transcription Activating RNA (STAR) encoding DNA templates are produced from viral genomic material within an RPA reaction. This reaction product is then added into a cell-free gene expression reaction for signal processing and visual output.

**Plant virus nucleic acid sources:** Cell lysates were prepared by grinding 20mg of *N. tabacum* leaf in lysis buffer (150mM KCl, 25mM Tris pH 7.5, 5mM EDTA, 5mM MgCl2, 0.5% NP-40, 1× HALT protease inhibitor cocktail (ThermoFisher Scientific), 0.5mM DTT, 100 U/ml RNase OUT (ThermoFisher Scientific)), and rapidly snap freezing in liquid nitrogen before storing at –80◦C. Cucumber mosaic virus infected material was similarly obtained from symptomatic *N. tabacum* leaf taken from plants inoculated, two weeks prior, with Bn57-CMV. PVY transcripts were generated from T7-tagged PCR products covering two regions of the PVY-O (Acc. No. AJ890349) genome (21-936nts and 6903-7899nts). CMV transcripts were generated from PCR product covering Bn57-CMV's 3$^{rd}$ RNA tagged with a T7 promoter. IVT reactions consisted of 10ul of 10X T7 Polymerase Buffer (NEB), 8ul of NTPs [25mM], 5ul DTT [100mM], 0.3 RNase Inhibitor [Promega], 2ul T7 RNA Polymerase (NEB), 20ul of the PCR product [~50nM], and 55ul H2O. Product was ethanol precipitated and run through a 20% Acrylamide gel with the correct band being excised with UV shadowing. This product was then eluted overnight in TE buffer and ethanol precipitated.

**RPA**: Recombinase Polymerase Amplification (RPA) reactions were prepared according to the manufacturer's protocol (Twist-DX TwistAMP® Liquid Basic RT). RPA reactions were prepared with 2.4µl of 10µM forward and reverse primer specific to the viral sample input (Supplementary Table 1), 29.5µl buffer, 11.2µl water, 2U Protector RNase Inhibitor (Roche), and 1µl of analyte (IVT product RNA at a concentration of 1nM, infected lysate, uninfected lysate, or water control) for a 50µl total volume. Reactions were run at 41$^{o}$C in a thermocycler without a heated lid for 8 minutes. Reactions were then gently mixed and microfuged followed by an additional 41$^{o}$C incubation for 32 minutes. 2.4ul of this product was utilized within each corresponding cell free reaction.

**Cell Free Extract Preparation:** Cell free extract and buffer were prepared as previously described with the buffer being supplemented with 80mM K-glutamate and 2mM Mg-Glutamate (1), and the cell lysis step performed with sonication. Sonication was performed with a sonicator (QSonica) using 6 pulses for 10 seconds with 10 second pauses in-between, at an amplitude of 50% (2).

**Cell Free Reactions:** 56.43µl of cell extract were mixed with 71.82µl of corresponding buffer. This mixture was supplemented with 2 mM Mg-Glutamate [60mM Stock], 6.67 mM Chi6 DNA [250mM Stock], 5.5 nM T7 expressing plasmid [610mM Stock], and 1 mM catechol [41mM Stock](Tokyo Chemical Industry) and then preincubated on a thermocycler at 29$^{o}$ C for 10 minutes. Following incubation, 7 nM of linearized target reporter plasmid was added along with 2.4 µl of the RPA reaction product. These target reporter plasmids were linearized with PCR and purified with PCR Purification Kit (Qiagen). 10 µl of the final reaction mix was placed into 3 separate wells on a 384 well microplate (Nunc Cat# 3712), sealed with an optically clear plate seal (Thermo Sci Cat#232701), and loaded into a SynergyH1 microplate reader (Biotek). The reader was programmed to observe absorbance at 385nm, sampling at 3 min intervals, while holding at 31$^{o}$C.

**Body Heat Amplification**: To test PLANT-Dx using body heat, an RPA reaction from CMV-infected or uninfected Arabidopsis plant lysate was setup as above. Reaction vessels were then taped under the axilla and pressed against the body and arm for 40 min of incubation. Following the incubation, 2.4 µl of the RPA reaction product was added into a cell free reaction containing CMV reporter constructs (Supplementary Table 1) prepared in the same manner as above, and tapped onto the lower forearm for 2 hours and 30 minutes.

**Limit Of Detection**: In vitro transcription was used to produce a representative 4nM stock of CMV's 3$^{rd}$ RNA. This stock was serially diluted in 1/10 by adding 5µl to 45µl of water down to 44fM. One µl from each dilution was then used in the PLANT-Dx procedure described above, and analyzed on the SynergyH1 microplate reader (Biotek).

**Photography:** All images were taken using an iPhone 8 (Apple) with the default photography application. The background of images was cropped using the masking tool within Adobe Illustrator.

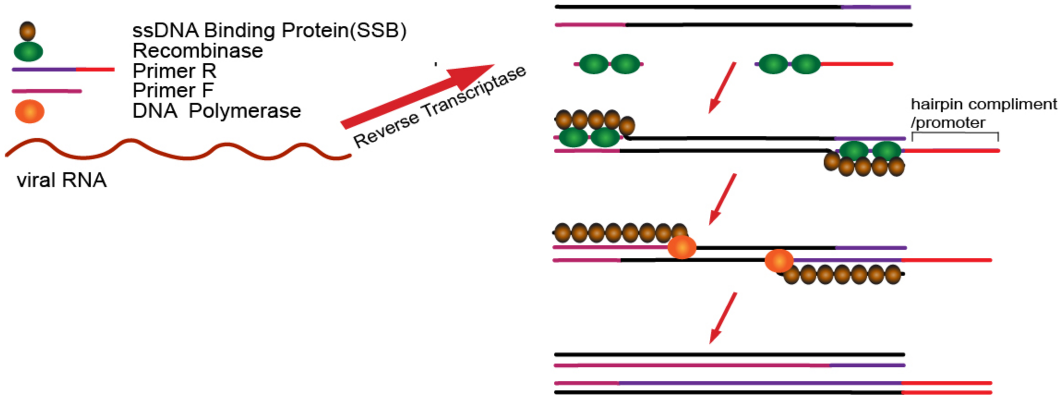


**Supplementary Figure 1:** Recombinase Polymerase Amplification (RPA) (3). Reverse transcriptase is utilized to create a cDNA construct from a target RNA genome. A recombinase facilitates the strand invasion and binding of designed primers to their target location within this construct. Single stranded DNA binding proteins can then stabilize the formation of a D loop enabling this primer invasion. Strand displacing polymerase can then extend the oligo to produce a dsDNA construct that can then template the next round for overall exponential amplification in an isothermal reaction. Overhanging sequence can be designed into the primers which are incorporated into the final dsDNA product. In this work overhang sequences are used to add a T7 RNA polymerase promoter and a fragment of a STAR sequence (Figure 1A) for downstream detection of an RNA produced from the RPA product.

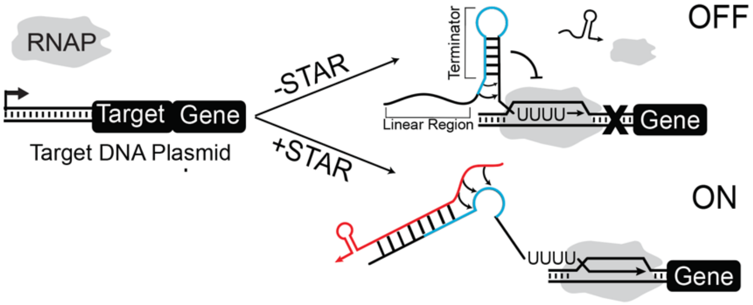


**Supplementary Figure 2:** Small Transcription Activating RNA (STAR) mechanism (4). Target DNA plasmid consists of a reporter gene transcriptionally gated by the presence of a 5' terminating hairpin (Target). In the absence of a STAR, the terminating hairpin folds, preventing downstream transcription of the reporter gene (OFF). When present, the STAR RNA binds to the linear region and 5' stem of the terminator, destabilizing the structure and allowing for downstream transcription of the desired reporter gene (ON).

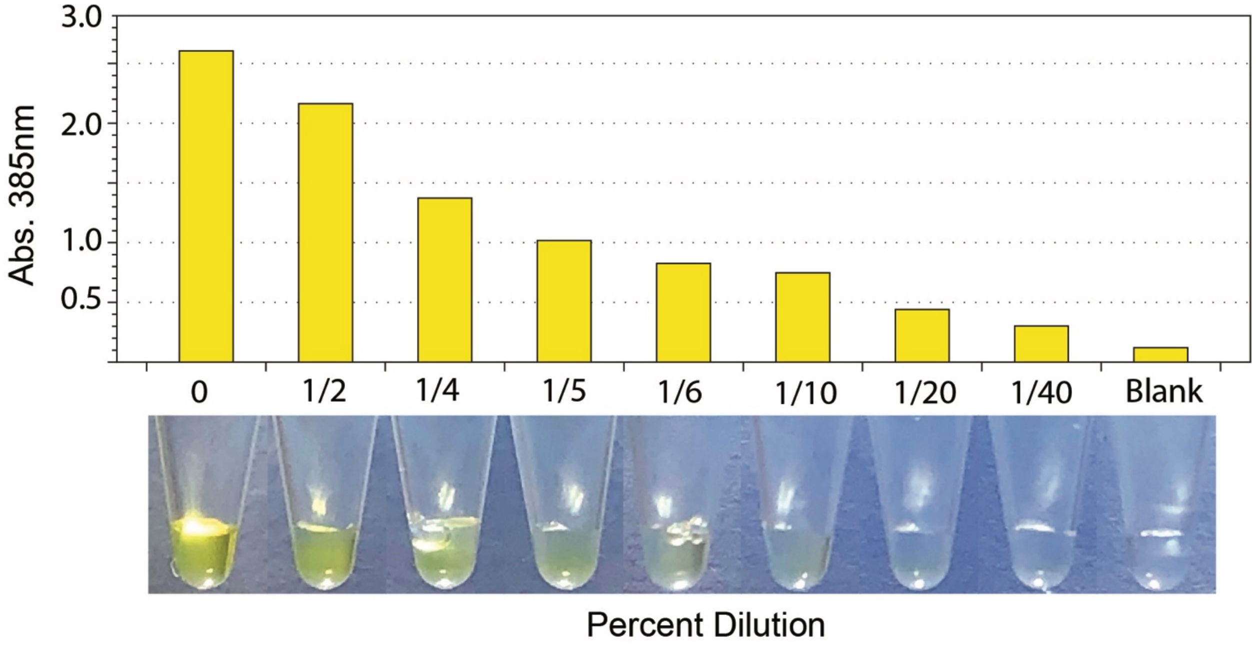


**Supplementary Figure 3:** Correlation between absorbance value on a SynergyH1 microplate reader (Biotek) with visual perception of color change in a catechol expression experiment. Reactions were allowed to proceed until a yellow pigment was clearly visible and then serially diluted with water. These dilutions were first photographed and then their absorbance was measured on the plate reader. In some instances, depending on the lighting and individual, it has been reported that it is possible to discern down to 0.5 Abs(385nm). Based upon this data, we have identified an absorbance of 0.8 to be our limit of quantitation (LOQ).

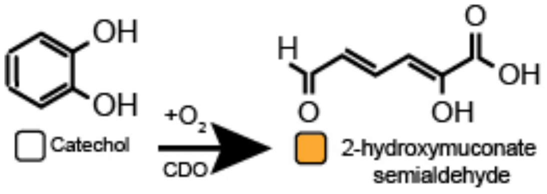


**Supplementary Figure 4:** Catechol to hydroxymuconic semialdehyde reaction. The enzyme catechol 2,3-dioxygenase (CDO) rapidly cleaves catechol into hydroxymuconic semialdehyde. This product can be detected by a measure of the absorbance at 385nm (Supplementary Figure 3).

**Supplementary Table 1:** Sequences of constructs and primers. Constructs for this work utilized nucleic acid sequences for flanking sequences (grey), promoters (blue), transcription terminating hairpins (red), ribosome binding sites (RBS) (purple), the coding sequence to our reporter gene CDO (gold), and the transcription terminator TrrnB (light blue).

| Name | Sequence 5' to 3' |
|---|---|
| Cucumber Mosaic Virus strain Bn57 Genomic Target | attaaccacccaacctttgtagggagtgagcgttgtaaacctggatacacgttcacatctat<br>taccctaaagccaccaaaa |
| RPA CMV Reverse Primer : Protective Sequence/T7 promoter/AD1 Hairpin antisense/CMV Target Region | gcatgacattaacctccagcaatctacatatcttaatacgactcactatagggtgaactgta<br>tacattccccgcttttggtggctttagggtaatagatg |
| CMV Forward Primer: Protective Sequence/ CMV Target Region | aaaaaaaacgccgcctttcggcggcgtttgattaaccacccaacctttgtagggagtgag |
| CMV Reporter Construct: Anderson Promoter J23119/ CMV Target Region/ AD1 Hairpin/ RBS / Catecholase coding sequence / TrrnB | ttgacagctagctcagtcctaggtataatactagtcacacgttcacatctattaccctaaag<br>ccaccaaaagcggggaatgtatacagttcatgtatatattccccgcttttttttttggatctagg<br>aggaaggatctatgaacaaaggtgtaatgcgaccgggccatgtgcagctgcgtgtactg<br>gacatgagcaaggccctggaacactacgtcgagttgctgggcctgatcgagatggacc<br>gtgacgaccagggccgtgtctatctgaaggcttggaccgaagtggataagttttccctggt<br>gctacgcgaggctgacgagccgggcatggattttatgggtttcaaggttgtggatgagga<br>tgctctccggcaactggagcgggatctgatggcatatggctgtgccgttgagcagctacc<br>cgcaggtgaactgaacagttgtggccggcgcgtgcgcttccaggccccctccgggcatc<br>acttcgagttgtatgcagacaaggaatatactggaaagtggggtttgaatgacgtcaatc<br>ccgaggcatggccgcgcgatctgaaaggtatggcggctgtgcgtttcgaccacgccctc<br>atgtatggcgacgaattgccggcgacctatgacctgttcaccaaggtgctcggtttctatct<br>ggccgaacaggtgctggacgaaaatggcacgcgcgtcgcccagtttctcagtctgtcga<br>ccaaggcccacgacgtggccttcattcaccatccggaaaaaggccgcctccatcatgtg<br>tccttccacctcgaaacctgggaagacttgcttcgcgccgccgacctgatctccatgacc<br>gacacatctatcgatatcggcccaacccgccacggcctcactcacggcaagaccatct<br>acttcttcgacccgtccggtaaccgcaacgaagtgttctgcgggggagattacaactacc<br>cggaccacaaaccggtgacctggaccaccgaccagctgggcaaggcgatcttttacca<br>cgaccgcattctcaacgaacgattcatgaccgtgctgacctgaggatctgaagcttgggc<br>ccgaacaaaaactcatctcagaagaggatctgaatagcgccgtcgaccatcatcatcat<br>catcattgagtttaaacggtctccagcttggctgttttggcggatgagagaagattttcagcc<br>tgatacagattaaatcagaacgcagaagcggtctgataaaacagaatttgcctggcggc<br>agtagcgcggtggtcccacctgaccccatgccgaactcagaagtgaaacgccgtagc<br>gccgatggtagtgtggggtctccccatgcgagagtagggaactgccaggcatcaaata<br>aaacgaaaggctcagtcgaaagactgggcctttcgttttatctgttgtttgtcggtgaact |
| Potato Virus Y Genomic Target | cgcattcagaagaaactggagaggaaggatagggaagaatatcacttccagatggcc<br>gctcctagtattgtgtcaaaa |
| RPA PVY Reverse Primer: Protective Sequence/T7 promoter/AD1 Hairpin | cccatgacattaactagcagcaatctacatatcttaatacgactcactatagggtgaactgt<br>atacattccccgcacgcattcagaagaaactggagagg |

| | |
|---|---|
| antisense/PVY target region | |
| RPA PVY Forward Primer: Protective Sequence/ CMV Target Region | aaaaaaaacggcgcctttcggcgccgtttgttttgacacaatactaggagcggccatctg |
| PVY Reporter Construct: Anderson Promoter J23119/ PVY Target Region/ AD1 Hairpin/ RBS / Catecholase coding sequence / TrrnB | ttgacagctagctcagtcctaggtataatactagtttcttccctatccttcctctccagtttcttctgaatgcgcgcggggaatgtatacagttcatgtatatattccccgcttttttttt ggatctaggaggaaggatctatgaacaaaggtgtaatgcgaccgggccatgtgcagctgcgtgtactggacatgagcaaggccctggaacactacgtcgagttgctgggcctgatcgagatggaccgtgacgaccagggccgtgtctatctgaaggcttggaccgaagtggataagttttccctggtgctacgcgaggctgacgagccgggcatggattttatgggtttcaaggttgtggatgaggatgctctccggcaactggagcgggatctgatggcatatggctgtgccgttgagcagctacccgcaggtgaactgaacagttgtggccggcgcgtgcgcttccaggccccctccgggcatcacttcgagttgtatgcagacaaggaatatactggaaagtggggtttgaatgacgtcaatcccgaggcatggccgcgcgatctgaaaggtatggcggctgtgcgtttcgaccacgccctcatgtatggcgacgaattgccggcgacctatgacctgttcaccaaggtgctcggtttctatctggccgaacaggtgctggacgaaaatggcacgcgcgtcgcccagtttctcagtctgtcgaccaaggcccacgacgtggccttcattcaccatccggaaaaaggccgcctccatcatgtgtccttccacctcgaaacctgggaagacttgcttcgcgccgccgacctgatctccatgaccgacacatctatcgatatcggcccaacccgccacggcctcactcacggcaagaccatctacttcttcgacccgtccggtaaccgcaacgaagtgttctgcgggggagattacaactacccggaccacaaaccggtgacctggaccaccgaccagctgggcaaggcgatcttttaccacgaccgcattctcaacgaacgattcatgaccgtgctgacctgaggatctgaagcttgggcccgaacaaaaactcatctcagaagaggatctgaatagcgccgtcgaccatcatcatcatcatcattgagtttaaacggtctccagcttggctgttttggcggatgagagaagattttcagcctgatacagattaaatcagaacgcagaagcggtctgataaaacagaatttgcctggcggcagtagcgcggtggtcccacctgaccccatgccgaactcagaagtgaaacgccgtagcgccgatggtagtgtggggtctccccatgcgagagtagggaactgccaggcatcaaataaaacgaaaggctcagtcgaaagactgggcctttcgttttatctgttgtttgtcggtgaact |

**Supplementary Note 1:** NUPACK script for discovering unstructured 40bp stretches from the Cucumber Mosaic Virus genome. Each reporter plasmid was designed with a novel linear region 5' of the terminating hairpin mimicking a 40bp stretch of the cognate pathogen of interest's genome. Building off of the design rules elucidated in Chappell et al. NUPACK was utilized to scan the genomes for regions with as little secondary structure as possible to ensure robust dynamic range from each construct.

```
material = rna
temperature = 37.0
trials = 3
structure line = U40
domain a = N40
source CMV =
GTAATCTTACCACTGTGTGTGTGCGTGTGTGTGTGTCGAGTCGTGTTGTCCGCACATTTGAGTCGTGTT
GTCCGCACATTTTATTTTTATCTTTGTACAGTGTGTTAGATTTCCCGAGGCATGGCTTTCCAAGGTACC
AGTAGGACTTTAACTCAACAGTCCTCAGCGGCTACGTCTGACGATCTTCAAAAGATATTATTTAGCCCT
GAAGCCATTAAGAAAATGGCTACTGAGTGTGACCTAGGCCGGCATCATTGGATGCGCGCTGATAATGCT
ATTTCAGTCCGGCCCCTCGTTCCCGAAGTAACCCACGGTCGTATTGCTTCCTTCTTTAAATCTGGATAT
GATGTTGGTGAATTGTGCTCAAAAGGATACATGAGCGTCCCTCAGGTGTTGTGTGCTGTTACTCGAACG
GTTTCCACCGATGCTGAAGGGTCTTTGAGAATTTACTTAGCTGATCTAGGTGACAAGGAGTTATCTCCT
ATAGATGGGCAATGCGTTTCGTTACATAACCATGATCTTCCCGCTTTGGTGTCTTTCCAACCGACGTAT
GACTGTCCTATGGAAACAGTTGGGAATCGCAAGCGGTGTTTTGCTGTCGTTATCGAAAGACATGGTTAC
ATTGGGTATACCGGTACCACAGCTAGCGTGTGTAGTAATTGGCAAGCAAGGTTTTCTTCTAAGAATAAC
AACTACACTCATATCGCAGCTGGGAAGACTCTAGTACTGCCTTTCAACAGATTAGCTGAGCAAACAAAA
CCGTCAGCTGTTGCTCGCCTGTTGAAGTCGCAATTGAACAACATTGAATCTTCGCAATATCTGTTAACG
AATGCGAAGATTAATCAGAATGCGCGCAGTGAGTCCGAGGAATTAAATGTTGAGAGCCCTCCCGCCGCA
ATCGGGAGTTCTTCCGCGTCCCGCTCCGAAGCCTTCAGACCGCAGGTGGTTAACGGTCTTTAGCACTTT
GGTGCGTATTAGTATATAAGTATTTGTGAGTCTGTACATAATACTATATCTATAGTGTCCTGTGTGAGT
TGATACAGTAGACATCTGTGACGCGATGCCGTGTTGAGAAGGGAACACATCTGGTTTTAGTAAGCCTAC
ATCACAGTTTTGAGGTTCAATTCCTCTTACTCCCTGTTGAGCCCCTTACTTTCTCATGGATGCTTCTCC
GCGAGATTGCgttat
window cmv_window = a
cmv_window.source = CMV
line.seq = a
prevent =  GGGG
```

**Supplementary References.**


1. Garamella, J. *et al*. The All E. Coli TX-TL Toolbox 2.0: A platform for Cell-Free Synthetic Biology. *ACS Syn. Bio.* 5(4) 344-55 (2016).
2. Silverman, A. *et al*. Deconstructing Cell-Free Extract Preparation for *In-Vitro* Activation of Transcriptional Genetic Circutry. *Biorxiv.* (2018). doi: 10.1101/411785
3. Piepenburg, O. et al. DNA Detection Using Recombination Proteins. *PLoS Biol.* 4(7) (2006).
4. Chappell, J., Westbrook, A., Verosloff, M., Lucks, J., Computational Design of Small Transcription Activating RNAs for Versatile and Dynamic Gene Regulation. *Nature Communications*. (2017). doi: 10.1038/s41467-017-01082-6
